# Ecological boundaries and constraints on viable eco-evolutionary pathways

**DOI:** 10.1101/2022.11.21.517427

**Authors:** Kyle E. Coblentz, John P. DeLong

## Abstract

Evolutionary dynamics are subject to constraints ranging from limitations on what is physically possible to limitations on the pathways that evolution can take. One set of evolutionary constraints, known as ‘demographic constraints’, constrain what can occur evolutionarily due to the population demographic or population dynamical consequences of evolution leading to conditions that make populations susceptible to extinction. These demographic constraints can limit the strength of selection or rates of environmental change populations can experience while remaining extant and the trait values a population can express. Here we further hypothesize that the population demographic and population dynamical consequences of evolution also can constrain the eco-evolutionary pathways that populations can traverse by defining ecological boundaries represented by areas of likely extinction. We illustrate this process using a model of predator evolution. Our results show that the populations that persist over time tend to be those whose eco-evolutionary dynamics have avoided ecological boundaries representing areas of likely extinction due to stochastic deviations from a deterministic eco-evolutionary expectation. We term this subset of persisting pathways viable eco-evolutionary pathways. The potential existence of ecological boundaries constraining evolutionary pathways has important implications for predicting evolutionary dynamics, interpreting past evolution, and understanding the role of stochasticity and ecological constraints on eco-evolutionary dynamics.

## Introduction

A diverse set of processes place constraints on evolution. These constraints occur on multiple levels from genetic constraints that limit evolutionary responses to selection to functional constraints that are due to limits on what is physically or physiologically possible (Arnold 1992, Kempes et al. 2012, 2019). Constraints also can limit the evolutionary pathways that evolution can traverse. For example, pathways in protein evolution can be limited to steps that retain protein function and evolution on genetic adaptive landscapes may have to occur in a manner that avoids ‘holes’ of low fitness on the landscape (Maynard Smith 1970, Gavrilets 1997, Poelwijk et al. 2007). The identification and understanding of these constraints on evolution is powerful because it allows us to narrow the domain over which evolutionary changes can occur.

An additional set of constraints on evolution are due to the consequences of evolution on population dynamics or population demography (Gomulkiewicz and Houle 2009, Amarasekare 2022). Population dynamic and demographic consequences of evolution are likely to constrain evolution because these can lead to situations in which populations become susceptible to extinction, even if the underlying genetics makes it physically possible for individuals to function. Therefore, these evolutionary dynamics are inaccessible for populations that are to remain persistent. For example, Gomulkiewicz and Houle (2009) have shown that populations are evolutionarily constrained because overly strong selection or rapid rates of environmental change are likely to lead to extinction. Similarly, a wide range of theoretical models have shown that natural selection can cause populations to evolve in a way that leads to their own extinction either deterministically or by causing populations to reach densities low enough to be susceptible to stochastic extinction in a process known as ‘Darwinian extinction’ or ‘evolutionary suicide’ (Matsuda and Abrams 1994, Webb 2003, Rankin and López-Sepulcre 2005, Parvinen and Dieckmann 2013). These studies suggest that populations that experience these sorts of selective forces without other sources of evolutionary constraints will not persist. Last, evolution may cause trait changes that lead to the destabilization of systems such as the development predator-prey cycles in which populations can become susceptible to stochastic extinction during the troughs of the cycles (Amarasekare 2022). Thus, evolutionary dynamics may be constrained away from these areas because populations with destabilizing traits are unlikely to persist over time. Altogether, these results suggest that, for populations that are to remain persistent, evolutionary dynamics may be constrained such that the demographic and population dynamic consequences of evolution do not lead to extinction.

Here we further this idea by developing the hypothesis that the demographic consequences of evolution also create ecological boundaries that constrain the eco-evolutionary pathways that populations can traverse. To illustrate this, imagine a population evolving on an adaptive landscape portraying fitness as a function of phenotypic trait values (Wright 1932, Simpson 1944, Svensson and Calsbeek 2013; Figure 1). Evolutionary theory predicts that the population will evolve towards peaks on the adaptive landscape following the path with the steepest fitness gradient (Figure 1A). However, evolution along the steepest fitness gradient may cause the population to reach trait values in which the extinction of the population is likely to occur (represented by the cut-out areas of the adaptive landscape in Figure 1B). This suggests that for populations that persist, evolution is more likely to proceed along trajectories that avoid these ecological boundaries through stochastic deviations from evolution along the steepest fitness gradient (Figure 1B). We term the resulting set of trajectories that avoid extinction viable eco-evolutionary pathways.

**Figure 1.**
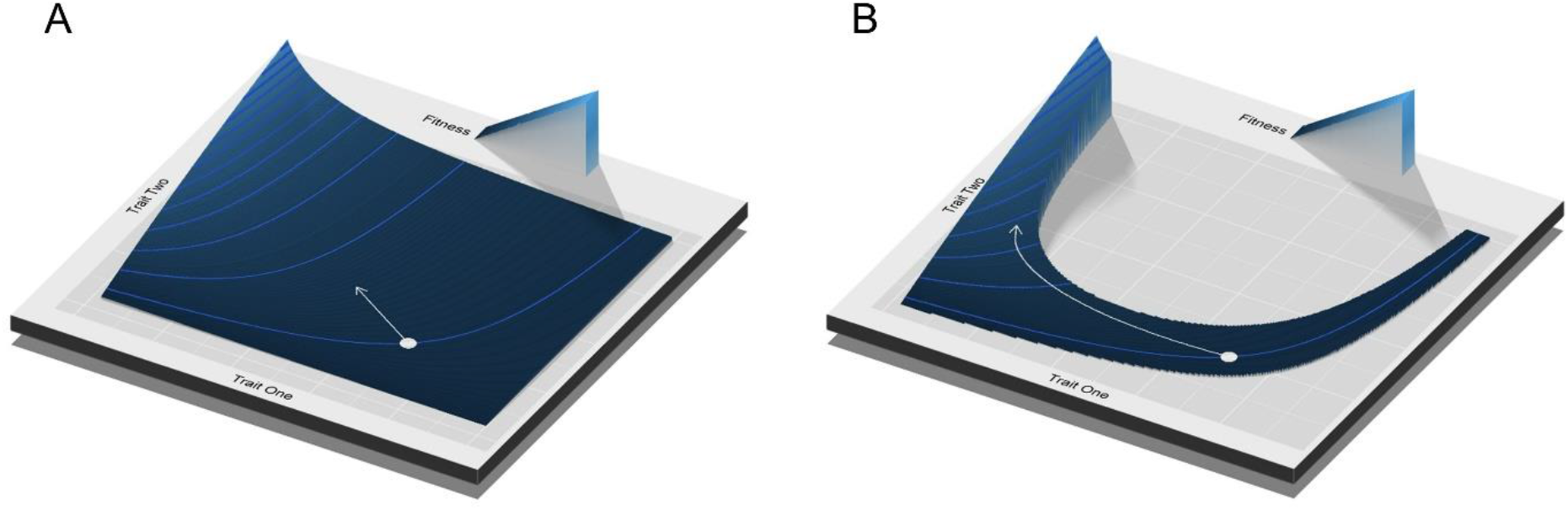
(A) Evolutionary theory predicts that populations will evolve towards higher fitness following the steepest fitness gradient. (B) However, some trait combinations may lead to ecological conditions in which the population is likely to go extinct, shown as areas cut out from the landscape. These boundaries will constrain persisting populations to a portion of the adaptive landscape, requiring their eco-evolutionary dynamics to follow only the viable eco-evolutionary pathways.

As a proof-of-concept that the demographic and population dynamic consequences of evolution can constrain viable eco-evolutionary pathways, and that populations may traverse these pathways by chance, we use a computational model of eco-evolutionary dynamics that directly incorporates demographic stochasticity and extinction. We specifically model predator feeding rate evolution in the widely used Rosenzweig-MacArthur predator prey model (Rosenzweig and MacArthur 1963). We use this model because the ecological properties of this model are well-known (e.g., what parameter values lead to stable vs. unstable dynamics and feasible vs. infeasible equilibria), the evolutionary expectations for the parameters governing the predator’s feeding rate (the space clearance (aka attack) rate and handling time) are well-known, and a recent study has shown that areas in which the dynamics of this model lead to predator-cycles are likely to cause extinction constraining the values that the predator’s feeding rate parameters can take (Rosenzweig and MacArthur 1963, Rosenzweig 1973, Johnson and Amarasekare 2015, Amarasekare 2022). In our analysis, we first use stochastic simulations in the absence of evolution to determine which areas of trait space are likely to lead to population dynamics that result in extinction. We then perform simulations in which the predator’s feeding rate parameters evolve to determine whether these identified regions of high extinction risk constrain the pathways that persistent populations can take. Overall, our results illustrate that the demographic and population dynamic consequences of evolution can constrain the viability of eco-evolutionary pathways.

## Methods

### Model

To examine the ability of ecological boundaries to constrain viable eco-evolutionary pathways, we analyze the evolution of a predator’s functional response parameters in the classic Rosenzweig-MacArthur predator-prey model (Rosenzweig and MacArthur 1963) in which the prey dynamics are described as:

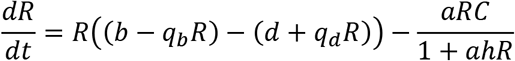

where *R* is the density of the prey, *b* is the birth rate of the prey, *d* is the natural mortality rate of the prey, *q_b_* and *q_d_* describe the density-dependence of the birth and natural mortality rates of the prey respectively, *a* is the predator’s space clearance rate of the prey, *h* is the predator’s handling time on the prey, and *C* is the predator density (parameter definitions are also given in Table 1). We explicitly model the prey’s birth and death rates and their density dependence to allow for a stochastic birth-death process and facilitate the use of the eco-evolutionary modeling approach we employ (see below). This form of logistic growth is equivalent to the classical model of logistic growth with intrinsic growth rate *r* = *b* – *d* and carrying capacity 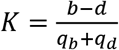 (DeLong and Coblentz 2022; See Supplemental Information S1). The predator’s dynamics are described as:

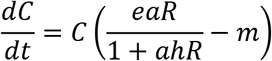

where *e* is the conversion efficiency of prey into predators, *m* is the per capita mortality rate of the predator, and all other parameters are defined above. This model has well-known stability and feasibility boundaries related to the predator’s functional response parameters (*a* and *h*, Murdoch *et al*., 2013; Johnson & Amarasekare, 2015). Specifically, the equilibrium of the model with positive densities of the predator and prey is unstable and leads to limit cycles when 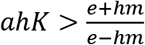 (Murdoch et al. 2013, Johnson and Amarasekare 2015). The predator cannot persist in the system when 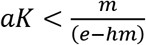 (Murdoch et al. 2013, Johnson and Amarasekare 2015). The evolution of the predator’s functional response parameters are also easily predicted as the selection gradient is always positive for *a* and negative for *h* (Rosenzweig 1973). That is, predator fitness increases with increasing space clearance rates and decreasing handling times.

**Table 1.** Parameters used in the models, their definitions, and the values taken in this study.

| Parameter | Definition | Values Used |
| --- | --- | --- |
| $b$ | Birth rate | 3.5 |
| $q_b$ | per capita density-dependent<br>decrease in the birth rate | 0.00875 when $K = 200$<br>0.000875 when $K = 2000$ |
| $d$ | Death rate | 1 |
| $q_d$ | Per capita density-dependent<br>increase in the death rate | 0.00875 when $K = 200$<br>0.000875 when $K = 2000$ |
| $K$ | Carrying capacity<br>$K = \frac{b - d}{q_b + q_d}$ | 200 or 2000 |
| $a$ | Space clearance rate | varied or evolves |
| $h$ | Handling time | varied or evolves |
| $e$ | Conversion efficiency | 0.3 when $K = 200$<br>0.1 when $K = 2000$ |
| $m$ | Per capita predator mortality<br>rate | 0.6 |
| $h_a^2$ | Narrow-sense heritability of the<br>space clearance rate in the<br>predator | 0.55 when $K = 200$<br>0.6 when $K = 2000$ |
| $h_h^2$ | Narrow sense heritability of the<br>handling time in the predator | 0.6 when $K = 200$<br>0.6 when $K = 2000$ |
| $CV_a$ | Coefficient of variation of the<br>initial log-normal distribution of<br>the predator space clearance rate | 0.33 when $K = 200$<br>0.8 when $K = 2000$ |
| $CV_h$ | Coefficient of variation of the<br>initial log-normal distribution of<br>the predator handling time | 0.325 when $K = 200$<br>0.4 when $K = 2000$ |

### GEMs and How They Work

To model predator evolution in the Rosenzweig-MacArthur model, we used Gillespie Eco-evolutionary Models (GEMs, (DeLong and Gibert 2016, Luhring and DeLong 2020, DeLong and Coblentz 2022, DeLong and Cressler in press)). GEMs work by adding an evolutionary component to the Gillespie algorithm for stochastic simulations of ordinary differential equations (ODEs; Gillespie, 1977). In short, GEMs are individual-based models that simulate differential equations and approximate the results of quantitative genetic analyses of phenotypic evolution while incorporating the effects of individual heterogeneity, demographic stochasticity, genetic drift, and the degradation of phenotypic variation with selection that are typically lacking from studies of eco-evolutionary dynamics. As stochasticity in evolutionary pathways and extinction are central to our hypothesis on ecological boundaries generating viable eco-evolutionary pathways, GEMs provide a useful tool for examining this hypothesis that would be difficult to examine using other tools for eco-evolutionary modelling.

Here we provide an in-depth explanation to how GEMs operate. First, the GEM is initiated with a matrix in which each row represents an individual from one of the considered populations (in the case here, either predators or prey). The columns of the matrix give the traits and parameters for each individual. For traits or parameters that evolve, the initial trait or parameter values for individuals are drawn from a log-normal distribution with a specified mean and coefficient of variation. All other traits or parameters receive the same value for all individuals within a population.

During each step of the GEM algorithm a random individual from each population is selected. The traits or parameters of these individuals are used to parameterize the ODE model underlying the GEM. The parameterized ODE is then broken up into corresponding ‘events’ as in the original Gillespie algorithm and which event occurs is determined randomly. For example, in our predator-prey model, the possible events are the birth of a prey, the natural death of a prey, the death of a prey via consumption by a predator, the birth of a predator, and the death of a predator. Which of these events occur during each iteration of the algorithm is randomly determined based on the relative magnitudes of the rates for each of the possible events. Specifically, we take a cumulative sum over all the possible events and then draw a random number determining which event occurs.

After determining which event occurs, that event is then played out through the modification of the matrix of individuals. In the case of a death in one of the populations, the individual selected from that population at the beginning of the step is removed from the population by deleting its corresponding row from the matrix. If a birth in a population occurs, a new row is added to the matrix of individuals. For traits or parameters that are not evolving, the values for these traits or parameters are simply placed in the corresponding column. For traits or parameters that are evolving, the new value of the trait is determined based on the value of the individual from the corresponding population selected at the beginning of the step and a chosen heritability of that trait or parameter using formulas derived from parent-offspring regression (DeLong and Belmaker 2019). Specifically, the new value is drawn from a log-normal distribution. The mean of the log-normal distribution is equal to 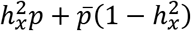 where 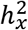 is the narrow-sense heritability of trait *x, p* is the trait of the parent, and 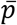 is the average of the trait in the population (DeLong and Belmaker 2019). The standard deviation of the log-normal distribution is equal to 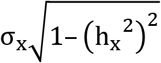 where *σ_x_* is the heritability-weighted mean of the initial and current standard deviations of the trait *x* or *σ_x_* = (1 – *h_x_*^2^)σ*_init_* + *h*_*x*_^2^σ_*current*_ (DeLong and Belmaker 2019). Thus, new individuals have a similar, but generally not identical, trait to their ‘parent’. Due to these methods of adding and removing individuals from populations, the algorithm performs a computational process analogous to natural selection with individuals with traits making them more ‘fit’ on average adding more similar individuals to the population on average compared to less ‘fit’ individuals. As in the original Gillespie algorithm, after an event resolves, the time is then advanced based on the expected time for the event to occur. This algorithm is repeated until a specified end time is reached with descriptions of population numbers and distributions of traits saved at standard times throughout the algorithm. Overall, this process then models the stochastic dynamics and trait evolution of populations based on a description of the system given by ODEs.

### Analysis Methods

Using GEMs, we first evaluated whether the analytical stability and feasibility boundaries of the model with respect to the predator’s functional response parameters defined a parameter space in which persistence was likely in the absence of evolution. To do so, we first simulated the dynamics of the predator and prey populations across a grid on the space clearance rate-handling time plane (*a*-*h* plane) on which the feasibility and stability boundaries of the system are defined. For each grid point on the plane, we performed 100 simulations over 50 time steps and calculated the proportion of simulations in which the predator or prey went extinct (i.e. reached a population size of zero). For the starting population values, we used the average population sizes following any transient dynamics from the deterministic solution. We also performed linear stability analysis of the deterministic model for each of the points on the *a*-*h* plane to determine the deterministic qualitative dynamics for that point (stable steady state, damped oscillations to a stable steady state, an unstable steady state leading to a stable limit cycle, or an infeasible steady state) and the resilience of the steady state equilibrium at that point measured by the maximum eigenvalue of the Jacobian matrix evaluated at the steady state (this eigenvalue determines the rate of return of the system to the equilibrium given a pulse perturbation away from the equilibrium; (McCann 2011, Murdoch et al. 2013)). For each point on the grid, we also determined the minimum population size of the deterministic dynamics after transient dynamics. In these simulations, no evolution occurred as we set the heritability of the space clearance rate and handling times and their variance within the population to zero. We did this for two carrying capacities (K = 200 and K = 2000) because higher carrying capacities lead to higher equilibrium population sizes which are likely to show different patterns of extinction due to demographic stochasticity (Giles Leigh 1981, Lande 1993, Ovaskainen and Meerson 2010).

After assessing the space in the *a-h* plane in which the predator and prey were likely to persist for each of the two carrying capacities in the absence of evolution, we then allowed predator populations’ space clearance rates and handling times to evolve directly and determined 1) whether the evolutionary pathways of populations that persisted differed from those of populations that went extinct and 2) whether extinction occurred in the areas of likely extinction identified in the absence of evolution. Although the selection gradients for the space clearance rate and handling time have constant signs, meaning that they are predicted to evolve to extreme and unrealistic values (infinity and zero, respectively), we are only interested in the trajectories of populations that persist over some time frame and the direct evolution of the functional response parameters is sufficient. Nevertheless, using an alternative model that assumes that the space clearance rate has a maximum, the handling time has a minimum value, and the predator has an evolving trait that determines both the space clearance rate and handling time gives similar answers to the model allowing the parameters to evolve directly (Supplemental Information S2). To match the simulations determining persistence on the *a-h* plane, these simulations also were run over 50 time steps. As we were particularly interested in whether stochastic eco-evolutionary dynamics could allow populations to take trajectories that avoided extinction, we specifically chose heritabilities and coefficients of variation for the space clearance rates and handling times that led to deterministic evolution to or near areas of likely extinction over the period of our simulations in quantitative genetics models.

Matlab and Mathematica code for the numerical analyses of the model are available (See Data Availability Statement).

## Results

### No Predator Evolution

The analytical stability and feasibility boundaries did largely define areas of likely population extinction with no evolution in the predator (Figures 2, 3). However, just considering these deterministic boundaries missed areas of likely extinction and persistence (Figures 2,3). For a carrying capacity of 200, an area of extinction occurred within the stability and feasibility boundaries at high space clearance rates and low handling times (Figure 2A). This area is associated with deterministic population dynamics that lead to damped oscillations to a steady state and stochastic dynamics that lead to quasi-cycles (cycles due to an interaction between the deterministic damped oscillations and stochasticity (Bartlett 1957, Gurney and Nisbet 1998, Pineda-Krch et al. 2007; Figures 2B). For areas in which the deterministic dynamics lead to damped oscillations and the stochastic dynamics lead to quasi-cycles, extinctions are more likely for the areas of the *a-h* plane that have lower resilience (a higher maximum eigenvalue; Figure 2C) and lower minimum population sizes of the deterministic dynamics (Figure 2D). For a carrying capacity of 2000, the area of extinction within the feasibility and stability boundaries at high space clearance rates and low handling times was reduced due to higher minimum population sizes and greater resilience despite the occurrence of quasi-cycles (Figure 3). Furthermore at a carrying capacity of 2000, there were some areas beyond but near the stability boundary where populations were able to persist despite dynamics leading to limit cycles of predator and prey abundances (Figure 3; note that the areas of persistence beyond the stability boundary at high handling times and space clearance rates occur because the simulation time of 50 time steps was shorter than the period of the limit cycles at these parameter values and all populations at these values would go extinct with longer simulation times).

**Figure 2.**
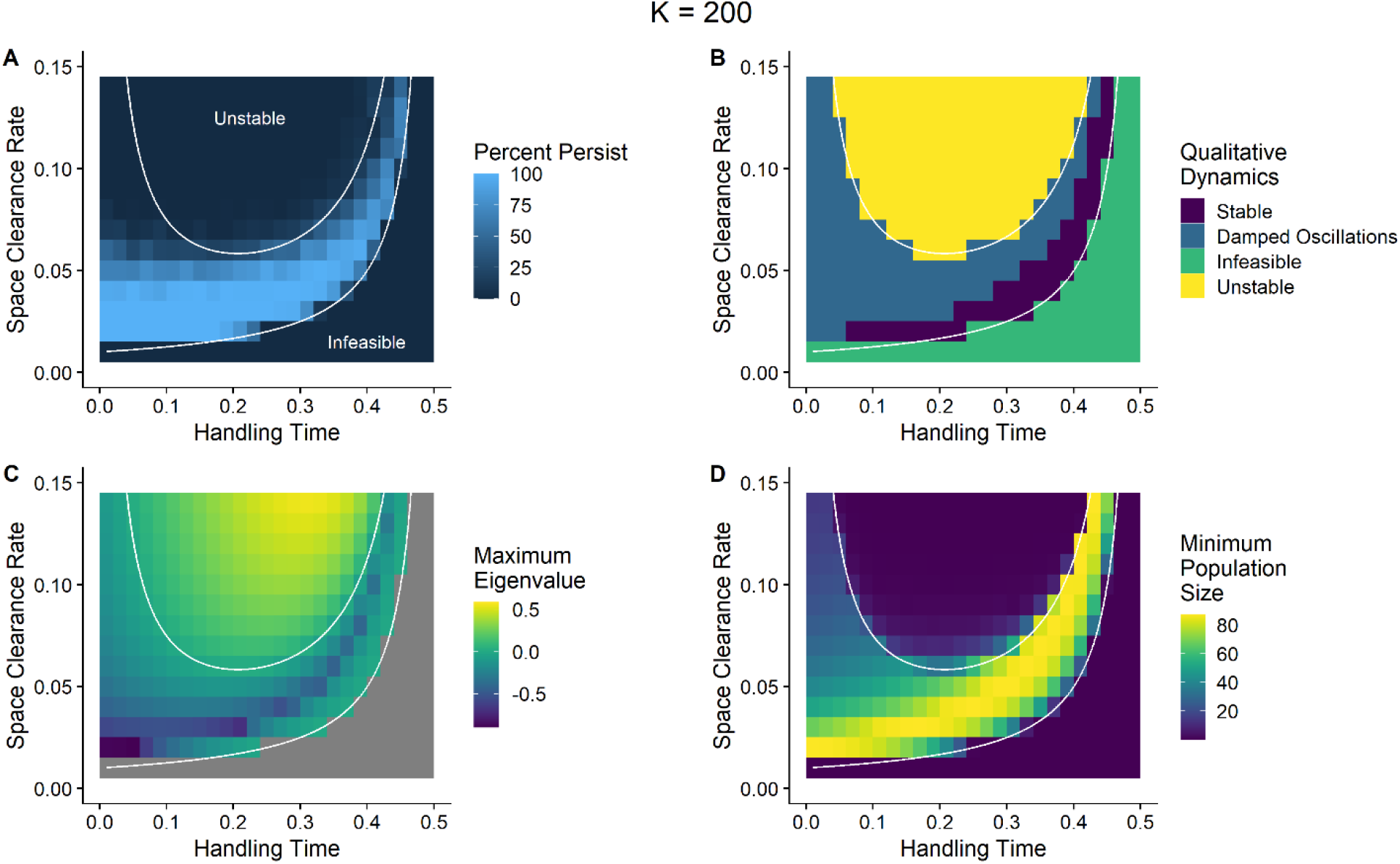
(A) In stochastic simulations of a predator-prey model, the percent of persistent populations varied within the stability and feasibility constraints predicted from the deterministic model (the white lines in Figures 2A-2D). Areas of extinction in the stochastic models occurred within the stability and feasibility boundaries. (B) Linear stability analysis of the deterministic models gives four areas with qualitatively different dynamics: a stable steady state (Stable), a stable steady state approached through damped oscillations (Damped Oscillations), an infeasible steady state with both predator and prey (Infeasible), and an unstable equilibrium leading to a stable limit cycle (Unstable). Areas within the stability and feasibility boundaries that lead to damped oscillations (B) but lead to persistent populations are associated with greater resilience (lower Maximum Eigenvalue, C) and higher minimum population sizes in deterministic dynamics (D). The non-a and -h parameters used are: b = 4.5, d = 1, q_b_ = q_d_ = 0.00875, e = 0.3, m = 0.6.

**Figure 3.**
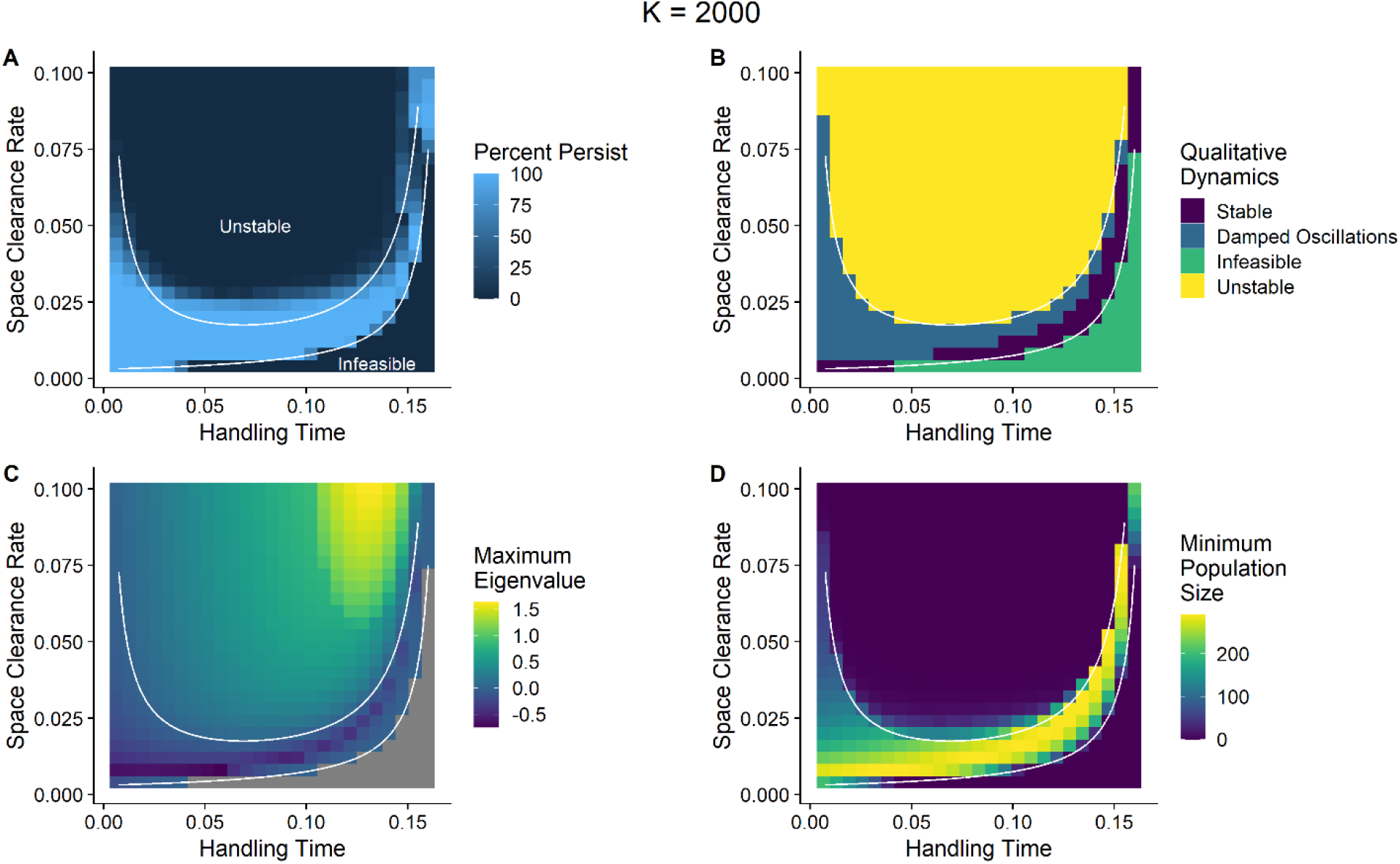
(A) In stochastic simulations of a predator-prey model, the percent of persistent populations varied within the stability and feasibility constraints predicted from the deterministic model (the white lines in Figures 2A-2D). Areas of extinction in the stochastic models occurred within the stability and feasibility boundaries and areas of persistence occurred beyond the stability boundary. (B) Linear stability analysis of the deterministic models gives four areas with qualitatively different dynamics: a stable steady state (Stable), a stable steady state approached through damped oscillations (Damped Oscillations), an infeasible steady state with both predator and prey (Infeasible), and an unstable steady state leading to a stable limit cycle (Unstable). Areas within the stability and feasibility boundaries that lead to damped oscillations (B) but lead to persistent populations are associated with greater resilience (lower Maximum Eigenvalue, C) and higher minimum population sizes in deterministic dynamics (D). The non-a and -h parameters used are: b = 4.5, d = 1, q_b_ = q_d_ = 0.000875, e = 0.1, m = 0.6.

### Predator Evolution

When predator populations evolved, populations that persisted evolved to parameter space that largely avoided the areas of likely extinction identified in the cases with no evolution (Figures 4, 5). In contrast, populations that went extinct tended to evolve higher space clearance rates at longer handling times causing the population to cross into areas of likely extinction (Figures 4A,B; 5A,B).

**Figure 4.**
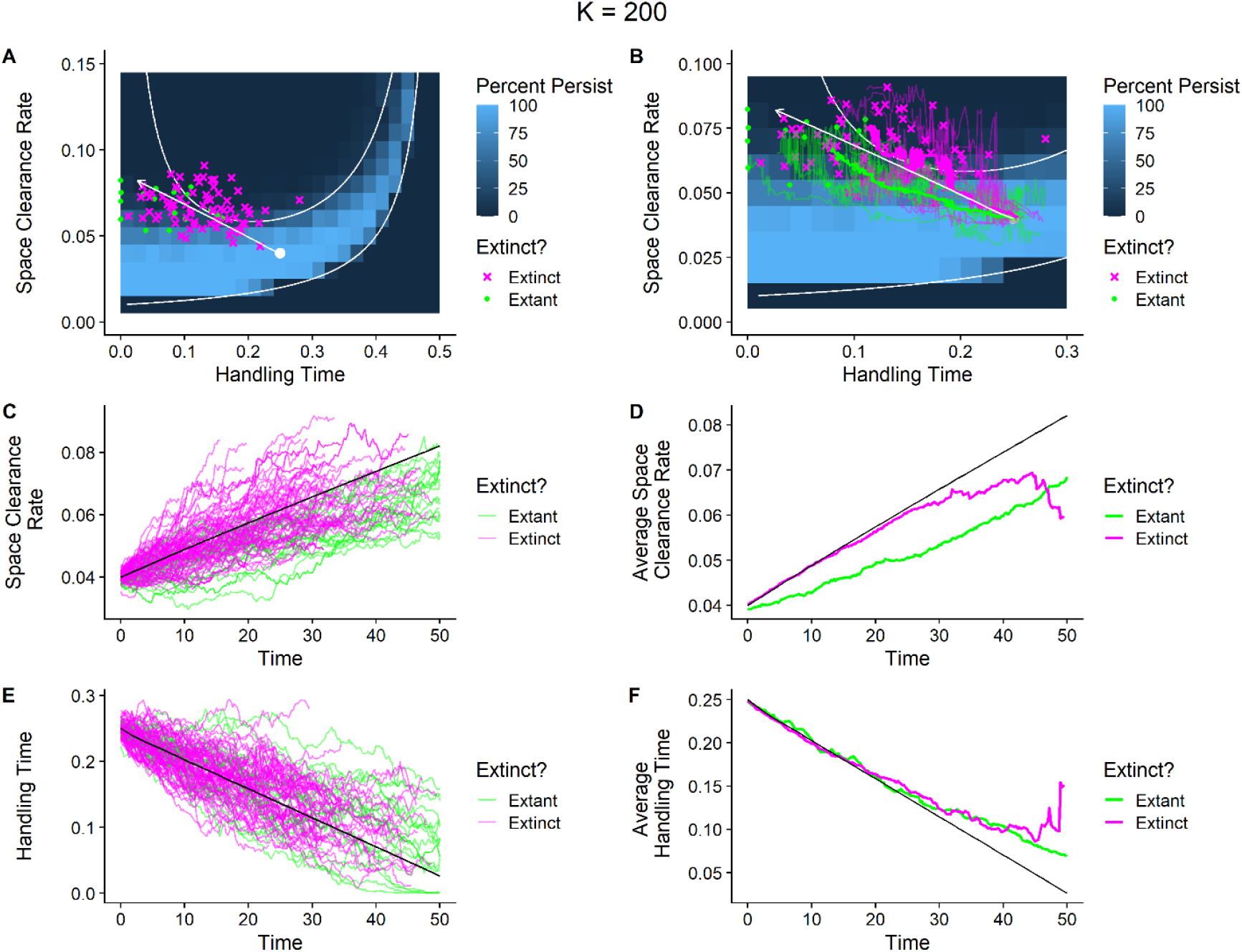
(A) Predator populations that persist (green points) after evolution tend to avoid areas of likely extinction, whereas predator populations that went extinct (magenta crosses) tended to evolve higher space clearance rates earlier at higher handling where extinction is more likely. The white arrow is the evolutionary pathway predicted by quantitative genetics. The white dot is the average starting values of the populations at the beginning of the simulation. (B,C,D) The evolutionary trajectories of space clearance rates for populations that went extinct (magenta lines) tended to reach higher space clearance rates earlier than for populations that were extant until the end of the simulations (green lines; thin lines in B represent the trajectories of 10 randomly chosen extinct and extant populations and the thick line is the average trajectory). (E,F) The evolutionary trajectories of handling times for populations that went extinct (magenta lines) showed no clear differences relative to populations that were extant and the end of the simulations (green lines). Black lines in C-F are the evolutionary trajectories predicted by quantitative genetics. The non-a and -h parameters used are: b = 4.5, d = 1, q_b_ = q_d_ = 0.00875, e = 0.3, m = 0.6. The heritability of a and h were 0.33 and 0.33 and the starting coefficients of variation in a and h among predators were 0.55 and 0.6, respectively.

**Figure 5.**
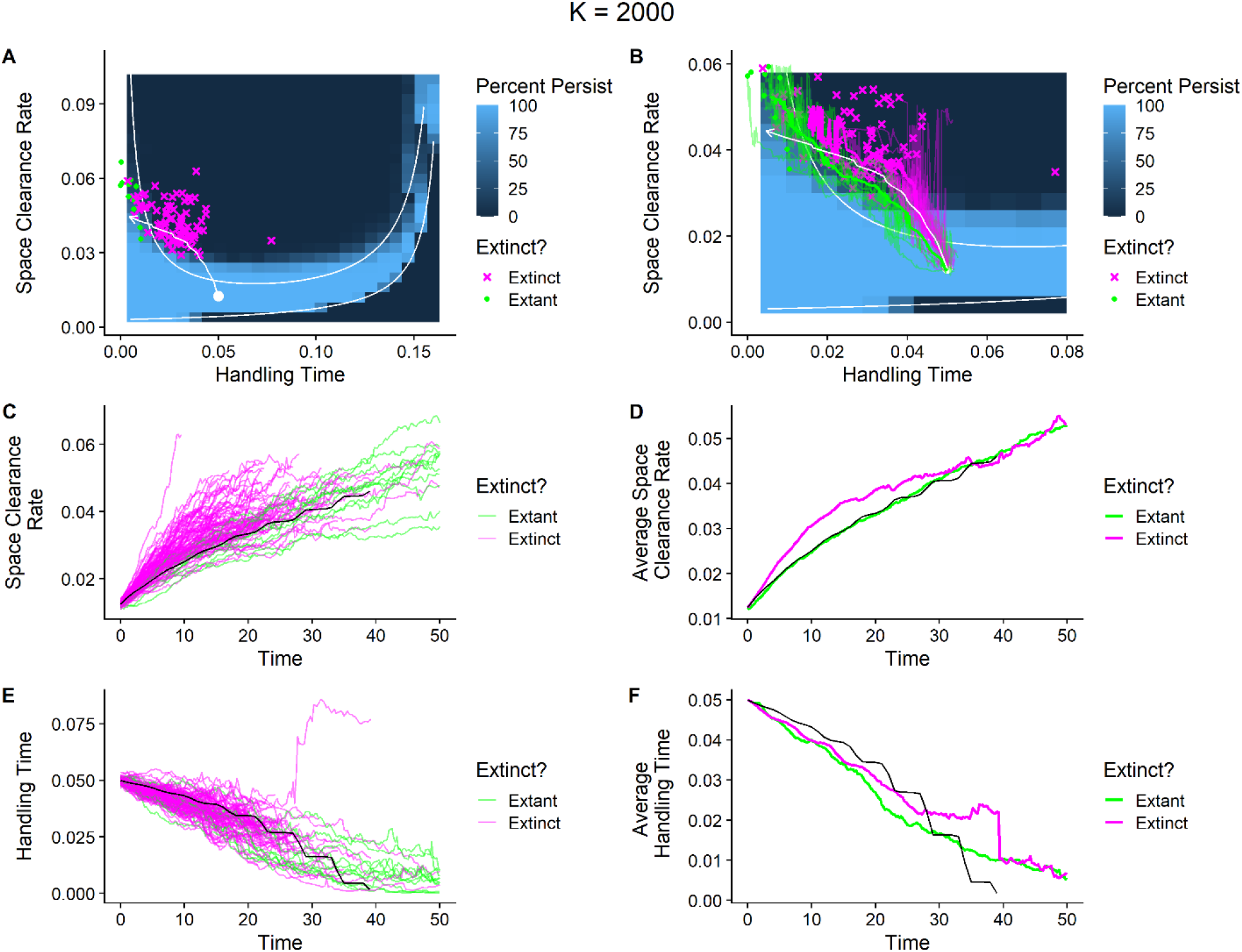
(A) Predator populations that persist (green points) after evolution tend to avoid areas of likely extinction, whereas predator populations that went extinct (magenta crosses) tended to evolve higher space clearance rates earlier into areas where extinction is more likely. The white arrow is the evolutionary pathway predicted by quantitative genetics. The white dot is the average starting values of the populations at the beginning of the simulation. (B, C, D) The evolutionary trajectories of space clearance rates for populations that went extinct (magenta lines) tended to reach higher space clearance rates earlier than for populations that were extant until the end of the simulations (green lines; thin lines represent the trajectories of 10 randomly chosen extinct and extant populations and the thick line is the average trajectory). (E,F) The evolutionary trajectories of handling times for populations that went extinct (magenta lines) evolved faster on average than populations that were extant and the end of the simulations (green lines). Black lines in C-F are evolutionary trajectories predicted by quantitative genetics. The non-a and -h parameters used are: b = 4.5, d = 1, q_b_ = q_d_ = 0.00875, e = 0.3, m = 0.6. The heritability of a and h were 0.33 and 0.33 and the starting coefficients of variation in a and h among predators were 0.55 and 0.6, respectively.

For a carrying capacity of 200 and populations that began with low space clearance rates and moderate handling times, the populations that persisted were those whose evolutionary dynamics stochastically led to slower evolution of the space clearance rate (Figures 4B-D). These populations on average showed a shallower evolutionary trajectory than that predicted by deterministic quantitative genetics (the white lines in Figures 4A,B and the black lines in Figures 4C-F). The evolutionary decrease in handling times of extant and extinct populations were similar and largely followed the quantitative genetics predictions but eventually slowed relative to the quantitative genetics prediction likely due to a decrease in trait variation over time (Figures 4E-F, Supplementary Information S3).

For a carrying capacity of 2000 and populations that began with low space clearance rates and moderate handling times, the populations that persisted were those whose evolutionary dynamics stochastically led to initially faster evolution of the handling time compared to extinct populations and the quantitative genetics prediction (Figures 5A-B, E-F). This initially fast decrease in the handling time kept evolutionary trajectories within areas where persistence was more likely (Figure 5B). In contrast, the evolutionary increase in the space clearance rate in the space clearance rate for persistent populations was similar to that predicted by quantitative genetics and generally slower than the populations that went extinct (Figures 5B-D).

## Discussion

Previous research has suggested that the population demographic and dynamical consequences of evolution can lead to constraints on the types of selection that species can experience and still persist (Gomulkiewicz and Houle 2009, Amarasekare 2022). Here using a stochastic, computational model of predator evolution, we show that areas that are likely to lead to population extinction establish ecological boundaries that define a set of viable eco-evolutionary pathways. This is because evolutionary dynamics that stochastically follow different trajectories can lead to two qualitatively different outcomes. First, some populations evolve trait combinations that lead to extinction by entering unstable, unfeasible, or low abundance ecological conditions susceptible to demographic stochasticity (i.e., Darwinian suicide). Second, populations that deterministically or by chance avoid ecological boundaries persist, following what become the viable subset of eco-evolutionary pathways.

Our results have implications for both forecasting evolutionary dynamics and interpreting past evolution. In terms of forecasting, our results illustrate the importance of considering the ecological consequences of evolutionary changes and incorporating the ecological environment into evolutionary forecasts (Nosil et al. 2020). Although a selection gradient may predict that a population evolve along a certain trajectory, if that trajectory leads to a trait space in which the ecological conditions are likely to lead to extinction, persistent populations may end up being those that have evolved along an alternative trajectory differing from that expected from the selection gradient. Operationalizing the existence of ecological barriers to predict actual empirical eco-evolutionary dynamics will require an increased connection between species’ traits and the parameters used in such models, although this may already be achievable in some laboratory eco-evolution systems (Yoshida et al. 2003, Kasada et al. 2014). Considering viable eco-evolutionary pathways is also important for interpreting past evolution. The persistence of a population suggests that that population’s evolution has avoided trait combinations likely to lead to that population’s extinction (Webb 2003, Borrelli et al. 2015). Thus, a population may not have evolved along the exact selection gradients occurring in the past and instead persisted because – possibly by chance alone – its evolutionary dynamics avoided ecological extinction boundaries. This suggests that past evolutionary dynamics may not always be informative about past selection gradients due to survivor bias (for other examples of survivor bias influencing evolutionary inference, see Budd and Mann (2018) and Weis (2018)). For example, if one observed the starting values of populations in Figure 4A and the ending values of the populations that persisted, they might infer a weaker selection gradient on predator space clearance rates than existed.

Stochasticity plays an important role in allowing populations to traverse viable eco-evolutionary pathways. First, because viable eco-evolutionary pathways may not match selection gradients, genetic drift is essential in allowing evolutionary dynamics to diverge from deterministic expectations. In our models, genetic drift is incorporated because individuals with traits or parameters that deterministically would lead to high fitness can stochastically die leaving few or no offspring (i.e. individual stochasticity *sensu* Caswell (2009)), while individuals with traits or parameters that would deterministically lead to low fitness may stochastically leave many offspring similar to themselves. Second, demographic stochasticity plays important roles in our models in both determining which areas of trait space represent high extinction risk and in generating variation among evolutionary trajectories. In terms of the areas that represent high extinction risk, our results show some mismatches between the predicted areas of high extinction risk from the deterministic model (i.e. the unstable and unfeasible areas of parameter space) and the areas of extinction that occurred in simulations with demographic stochasticity. At low carrying capacities, some of the areas of extinction not predicted by the deterministic model involved the presence of quasi-cycles that are the product of the interaction of damped oscillations in the deterministic model and demographic stochasticity and cannot occur without demographic stochasticity (Bartlett 1957, Gurney and Nisbet 1998, Pineda-Krch et al. 2007). At high carrying capacities, the simulations without evolution suggested that some parameter combinations that lead to predator-prey cycles may nevertheless still lead to persistent populations because populations do not reach low enough sizes to be susceptible to extinction via demographic stochasticity. In terms of the effects of demographic stochasticity in generating variation in evolution trajectories among populations, this is because the selection gradients on the predator’s space clearance rates and handling times depend on the densities of the prey (Rosenzweig 1973, Amarasekare 2022, DeLong and Coblentz 2022). Because demographic stochasticity can alter population abundances relative to deterministic expectations, two predator populations with exactly the same distribution of traits could experience different selection gradients because their prey population abundances are stochastically different. Altogether, the importance of stochasticity in generating our results highlights the importance of including stochasticity into theory on eco-evolutionary dynamics (Benaïm and Schreiber 2019, DeLong and Cressler in press).

Although our results highlight the role of stochasticity in allowing populations to traverse viable eco-evolutionary pathways, evolutionary constraints on traits also may operate to prevent species from evolving to areas in which ecological conditions are likely to lead to extinction. For example, counteracting selection from other sources or a lack of heritable variation may operate to prevent species from evolving into ecological scenarios likely to cause extinction (Arnold 1992, Vuorinen et al. 2021). In fact, if our simulations were run long enough eventually all populations would end up extinct. Thus, the existence of factors slowing or counteracting evolution are likely important and could possibly lead to a form of selection bias across populations in which populations that do exhibit evolutionary constraints that prevent their evolution to ecologically risky trait spaces are more likely to be observed (Webb 2003, Rankin and López-Sepulcre 2005, Parvinen and Dieckmann 2013). Note, however, that viable eco-evolutionary pathways also can occur in models in which the predator and prey will not evolve to extinction (Supplemental Information S2).

Despite our use of a predator-prey model to illustrate the concept of viable eco-evolutionary pathways, this process should be general to systems in which certain trait combinations lead to high likelihoods of extinction (e.g. low population sizes). We view this work as analogous to past research on limits to evolutionary trajectories. For example, Gavrilets (1997) suggested that adaptive landscapes representing the fitness of genotypic frequencies contain genotypes of low fitness (or inviability) creating ‘holes’ in the fitness landscape in which evolutionary trajectories must evolve around. Similarly, our results suggest that eco-evolutionary trajectories must avoid certain trait combinations that lead to ecological conditions with high extinction risk even while being genetically viable. Studies on protein evolution also have shown that amino acid substitutions may have to occur in a certain order for the evolving protein to remain functional or provide a fitness benefit (Maynard Smith 1970, Poelwijk et al. 2007). Similarly, ecological barriers may require that certain traits evolve before others for the population to follow the viable eco-evolutionary pathway. For example, in the scenario with evolution and a high carrying capacity, populations that evolved lower handling times quickly at the beginning of the simulation were able to achieve higher space clearance rates later in time while still persisting. In general, we expect that the ecological consequences of evolutionary changes along with intrinsic evolutionary constraints act to limit the evolutionary pathways persistent lineages can traverse.

## Literature Cited

Amarasekare, P. 2022. Ecological Constraints on the Evolution of Consumer Functional Responses. -Frontiers in Ecology and Evolution in press.

Arnold, S. J. 1992. Constraints on Phenotypic Evolution. -The American Naturalist 140: S85–S107.

Bartlett, M. S. 1957. Measles Periodicity and Community Size. -Journal of the Royal Statistical Society. Series A (General) 120: 48–70.

Benaïm, M. and Schreiber, S. J. 2019. Persistence and extinction for stochastic ecological models with internal and external variables. -J. Math. Biol. 79: 393–431.

Borrelli, J. J. et al. 2015. Selection on stability across ecological scales. -Trends in Ecology & Evolution 30:417–425.

Budd, G. E. and Mann, R. P. 2018. History is written by the victors: The effect of the push of the past on the fossil record. -Evolution 72: 2276–2291.

Caswell, H. 2009. Stage, age and individual stochasticity in demography. -Oikos 118: 1763–1782.

DeLong, J. P. and Gibert, J. P. 2016. Gillespie eco-evolutionary models (GEMs) reveal the role of heritable trait variation in eco-evolutionary dynamics. -Ecology and Evolution 6: 935–945.

DeLong, J. P. and Belmaker, J. 2019. Ecological pleiotropy and indirect effects alter the potential for evolutionary rescue. -Evolutionary Applications 12: 636–654.

DeLong, J. P. and Coblentz, K. E. 2022. Prey diversity constrains the adaptive potential of predator foraging traits. -Oikos in press.

DeLong, J. P. and Cressler, C. E. Stochasticity directs adaptive evolution toward nonequilibrium evolutionary attractors. -Ecology n/a: e3873.

Gavrilets, S. 1997. Evolution and speciation on holey adaptive landscapes. -Trends in Ecology & Evolution 12: 307–312.

Giles Leigh, E. 1981. The average lifetime of a population in a varying environment. -Journal of Theoretical Biology 90: 213–239.

Gillespie, D. T. 1977. Exact stochastic simulation of coupled chemical reactions. -J. Phys. Chem. 81: 2340–2361.

Gomulkiewicz, R. and Houle, D. 2009. Demographic and Genetic Constraints on Evolution. -The American Naturalist 174: E218–E229.

Gurney, W. and Nisbet, R. M. 1998. Ecological dynamics. -Oxford University Press.

Johnson, C. A. and Amarasekare, P. 2015. A Metric for Quantifying the Oscillatory Tendency of Consumer-Resource Interactions. -The American Naturalist 185: 87–99.

Kasada, M. et al. 2014. Form of an evolutionary tradeoff affects eco-evolutionary dynamics in a predator–prey system. -PNAS 111: 16035–16040.

Kempes, C. P. et al. 2012. Growth, metabolic partitioning, and the size of microorganisms. -Proceedings of the National Academy of Sciences 109: 495–500.

Kempes, C. P. et al. 2019. The Scales That Limit: The Physical Boundaries of Evolution. -Frontiers in Ecology and Evolution in press.

Lande, R. 1993. Risks of Population Extinction from Demographic and Environmental Stochasticity and Random Catastrophes. -The American Naturalist 142: 911–927.

Luhring, T. M. and DeLong, J. P. 2020. Trophic cascades alter eco-evolutionary dynamics and body size evolution. -Proc. R. Soc. B. 287: 20200526.

Matsuda, H. and Abrams, P. A. 1994. Runaway Evolution to Self-Extinction Under Asymmetrical Competition. -Evolution 48: 1764–1772.

Maynard Smith, J. 1970. Natural Selection and the Concept of a Protein Space. -Nature 225: 563–564.

McCann, K. S. 2011. Food Webs (MPB-50). -Princeton University Press.

Murdoch, W. W. et al. 2013. Consumer-Resource Dynamics (MPB-36). -Princeton University Press.

Nosil, P. et al. 2020. Increasing our ability to predict contemporary evolution. -Nat Commun 11: 5592.

Ovaskainen, O. and Meerson, B. 2010. Stochastic models of population extinction. -Trends in Ecology & Evolution 25: 643–652.

Parvinen, K. and Dieckmann, U. 2013. Self-extinction through optimizing selection. -Journal of Theoretical Biology 333: 1–9.

Pineda-Krch, M. et al. 2007. A Tale of Two Cycles: Distinguishing Quasi-Cycles and Limit Cycles in Finite Predator-Prey Populations. -Oikos 116: 53–64.

Poelwijk, F. J. et al. 2007. Empirical fitness landscapes reveal accessible evolutionary paths. -Nature 445: 383–386.

Rankin, D. J. and López-Sepulcre, A. 2005. Can adaptation lead to extinction? -Oikos 111: 616–619.

Rosenzweig, M. L. 1973. Evolution of the Predator Isocline. -Evolution 27: 84–94.

Rosenzweig, M. L. and MacArthur, R. H. 1963. Graphical Representation and Stability Conditions of Predator-Prey Interactions. -The American Naturalist 97: 209–223.

Simpson, G. G. 1944. Tempo and Mode in Evolution. -Columbia University Press.

2013. The Adaptive Landscape in Evolutionary Biology (E Svensson and R Calsbeek, Eds.). -Oxford University Press.

Vuorinen, K. E. M. et al. 2021. Why don’t all species overexploit? -Oikos 130: 1835–1848.

Webb, C. 2003. A Complete Classification of Darwinian Extinction in Ecological Interactions. -The American Naturalist 161: 181–205.

Weis, A. E. 2018. Detecting the “invisible fraction” bias in resurrection experiments. -Evolutionary Applications 11: 88–95.

Wright, S. 1932. The roles of mutation, inbreeding, crossbreeding and selection in evolution. -Proceedings of the Sixth Annual Congress of Genetics 1: 356–366.

Yoshida, T. et al. 2003. Rapid evolution drives ecological dynamics in a predator–prey system. -Nature 424: 303–306.

