## Supplemental Information for "Ecological boundaries and constraints on viable eco-evolutionary pathways"

**Contents** **Page**

**S1 Equivalence of Logistic Models** **2**

**S2 Alternative Model with Maximum Space Clearance Rate and Minimum Handling Time** **3**

**S3 Supplemental Figures** **7**

### S1 Equivalence of Logistic Models

Here we show the equivalence of the usual logistic growth equation

$$\frac{dN}{dt} = rN\left(1 - \frac{N}{K}\right) \qquad \text{eqn. S1.1}$$

where $N$ is the population size, $r$ is the intrinsic growth rate, and $K$ is the carrying capacity, with the birth/death formulation of the logistic growth we use in the main text

$$\frac{dN}{dt} = N[(b - q_bN) - (d + q_dN)] \qquad \text{eqn. S1.2}$$

where $N$ is the population size, $b$ is the birth rate, $q_b$ is the density dependence of the birth rate, $d$ is the death rate, and $q_d$ is the density dependence of the death rate.

We start with the right-hand side of equation S1.2

$$N[(b - q_bN) - (d + q_dN)] \qquad \text{eqn. S1.3}$$

Multiplying through by $N$ gives

$$= Nb - q_bN^2 - dN - q_dN^2 \qquad \text{eqn. S1.4}$$

Bringing together the $b$ and $d$ terms gives

$$= N(b - d) - q_bN^2 - q_dN^2 \qquad \text{eqn. S1.5}$$

We then factor $N(b - d)$ from each term to give

$$= N(b - d)[1 - \frac{q_bN}{(b-d)} - \frac{q_dN}{(b-d)}] \qquad \text{eqn. S1.6}$$

Last, we combine the final two terms to get

$$= N(b - d)\left[1 - \frac{(q_b+q_d)N}{(b-d)}\right] \qquad \text{eqn. S1.7}$$

Letting $r = (b - d)$ and $K = \frac{(b-d)}{q_b+q_d}$, one can see that the two models of logistic growth are equivalent.

### S2 Alternative Model with Maximum Space Clearance Rate and Minimum Handling Time

In the main text, we analyze a model of predator evolution to illustrate how areas of likely extinction can generate viable eco-evolutionary pathways. Although the model we use illustrates viable eco-evolutionary pathways, it has some unrealistic features. One of these features is that, eventually, any predator will evolve to extinction because the predator will evolve very high space clearance rates and very low handling times deterministically driving the prey and, hence, itself to extinction. Here we analyze an alternative model similar to that analyzed within Amarasekare (2022). In this model, rather than evolving ever higher space clearance rates and ever lower handling times, predators instead evolve towards a maximum space clearance rate and a minimum handling time. This mimics the case in which space clearance rates and handling times are determined by both predator and prey traits, but only the predator is evolving and can only achieve a maximum space clearance rate and minimum handling time set by the prey's traits (Amarasekare 2022).We show that this model gives similar qualitative results as we present in the main text. That is, populations that persist for the duration of the simulation are those whose eco-evolutionary trajectories generally avoid parameter combinations likely to lead to extinction.

In this model with a maximum space clearance rate and minimum handling time, we assume that predator individuals have two traits, $x_a$ and $x_h$, that determine their space clearance rates and handling times, respectively. We then assume that the values of these traits are related to the space clearance rate and handling time by unimodal functions with a maximum and minimum, respectively. By doing so, we assume that there is stabilizing selection for the predator's traits to evolve to reach the trait values that lead to the maximum space clearance rate and minimum handling time. The function translating the trait - $x_a$ into the space clearance is

$$a(x_a) = a_{\max} e^{-\frac{(\theta_a - x_a)^2}{2\tau_a{}^2}}, \qquad \text{eqn. S2.1}$$

where $a(x_a)$ is the realized space clearance rate of a predator with trait $x_a$, $a_{max}$ is the maximum space clearance rate, $\theta_a$ is the value of the trait $x_a$ at which the space clearance rate reaches its maximum, and $\tau_a$ describes the rate at which the space clearance rate decreases from the maximum as the trait $x_a$ moves away from $\theta_a$ (the definitions of the all of the parameters used in this supplemental material and the values that were used in the subsequent simulation are given in Table S2.1). The function translating the handling time trait $x_h$ into the handling time is an analogous function to that in eqn. S2.1 that has been flipped over the x-axis and translated upwards such that it has a minimum value that is greater than zero rather than a maximum value. The specific function we use is

$$h(x_h) = 1 + h_{min} - e^{-\frac{(\theta_h - x_h)^2}{2\tau_h^2}}, \qquad \text{eqn. S2.2}$$

where $h(x_h)$ is the realized handling time of a predator with trait $x_h$, $h_{min}$ is the minimum handling time, $\theta_h$ is the value of the trait $x_h$ at which the handling time reaches its minimum, and $\tau_h$ describes the rate at which the handling time increases from the minimum value as the trait $x_h$ moves away from $\theta_h$.

Given the functions translating between the predator traits and the space clearance rates and handling times, we model the predator and prey population dynamics using the Rosenzweig-MacArthur predator-prey model as in the main text. Specifically, the dynamics of the prey density $R$ are given by

$$\frac{dR}{dt} = R\big((b - q_b R) - (d - q_d R)\big) - \frac{a(x_a)RC}{a(x_a)h(x_h)R}, \qquad \text{eqn. S2.3}$$

where $b$ is the prey's birth rate, $d$ is the prey's death rate, and $q_b$ and $q_d$ are the density dependences of the birth and death rates, respectively, with all other parameters defined above. The dynamics of the predator density are given by

$$\frac{dC}{dt} = \frac{ea(x_a)RC}{1+a(x_a)h(x_h)R} - mC, \qquad \text{eqn. S2.4}$$

where $e$ is the conversion efficiency of prey into predators, and $m$ is the density independent mortality rate of the consumer with all other parameters defined above.

Given the model for the densities of the resource and the consumer, we analyzed the eco-evolutionary dynamics generated by the evolution in the predator of the traits determining the space clearance rate and handling time using GEMs as in the main text. Specifically, we used a similar parameter set as the high carrying capacity case in the main text and allowed both the trait determining the space clearance rate and the trait determining the handling time to evolve giving them an initial coefficient of variation ($CV_{x_a}$ and $CV_{x_h}$) and making them heritable (with heritabilities $h^2_{x_a}$ and $h^2_{x_h}$).

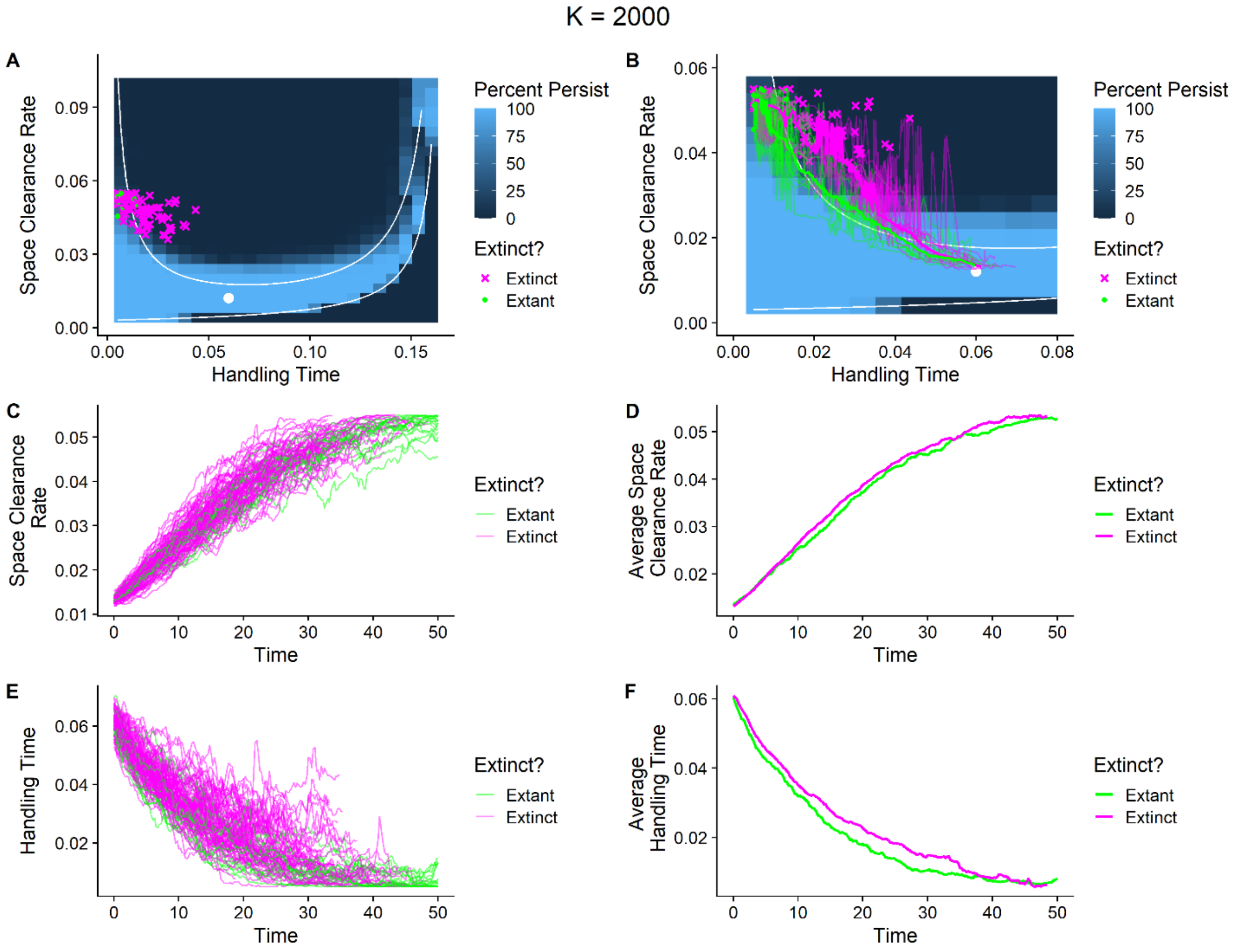


**Figure S2.1.** *(A) Predator populations that persist (green points) after evolution tend to avoid areas of likely extinction, whereas predator populations that went extinct (magenta crosses) tended to evolve higher space clearance rates earlier into areas where extinction is more likely. The white dot is the*

*average starting values of the populations at the beginning of the simulation. (A,B) The evolutionary trajectories of space clearance rates for populations that went extinct (magenta lines) tended increase more quickly than for populations that were extant until the end of the simulations (green lines; thin lines represent the trajectories of 10 randomly chosen extinct and extant populations and the thick line is the average trajectory). (E,F) The evolutionary trajectories of handling times for populations that went extinct (magenta lines) evolved slower on average than populations that were extant and the end of the simulations (green lines). The parameters used are:* $a_{max} = 0.05, h_{min} = 0.005, \theta_a = 8, \theta_h = 8, \tau_a = 10, \tau_h = 10, b = 3.5, q_b = 0.000875, d = 1, q_d = 0.000875, e = 0.1, m = 0.6, CV_{x_a} = 0.125, CV_{x_h} = 0.125, h^2_{x_a} = 0.2, h^2_{x_h} = 0.2.$

Using the model with a maximum space clearance rate and minimum handling time that the predator can achieve, we find that the predator populations that remain extant for the duration of the eco-evolutionary simulations are again those that largely avoided identified areas of likely extinction (Figure S2.1). In this model, this occurs as the predator populations evolve toward the parameter space given by the maximum achievable space clearance rate and minimum handling time. In particular, the extant populations in these simulations tend to be those that stochastically evolve lower handling times more quickly than the extinct populations and space clearance rates more slowly than the extinct populations on average (Figure S2.1). Overall, these results show that our conclusions regarding viable eco-evolutionary pathways are robust to the unrealistic feature of our model in the main text that predators can evolve arbitrarily high space clearance rates and arbitrarily small handling times.

**Table S2.1.** A table of the parameters used in the model presented in Supplemental Information S2, their definitions, and the values that we used for those parameters.

| Parameter | Definition | Value |
|---|---|---|
| $x_a$ | The trait value of the predator that determines the space clearance rate | Evolves |
| $x_h$ | The trait value of the predator that determines the handling time | Evolves |
| $a_{max}$ | Maximum value of the space clearance rate | 0.05 |
| $h_{min}$ | Minimum value of the handling time | 0.005 |
| $\theta_a$ | The value of $x_a$ at which the space clearance rate is maximized | 8 |
| $\theta_h$ | The value of $x_h$ at which the handling time is minimized | 8 |
| $\tau_a$ | Parameter determining the rate at which the space clearance rate declines from its maximum as $x_a$ moves away from $\theta_a$ | 10 |
| $\tau_h$ | Parameter determining the rate at which the handling time increases from its minimum as $x_h$ moves away from $\theta_h$ | 10 |
| $b$ | Birth rate of the prey | 4.5 |
| $q_b$ | Density dependence of the birth rate of the prey | 0.000875 |
| $d$ | Natural death rate of the prey | 1 |
| $q_d$ | Density dependence of the natural death rate of the prey | 0.000875 |
| $e$ | Conversion efficiency of prey into predators | 0.1 |
| $m$ | Density independent mortality of the predator | 0.6 |
| $CV_{x_a}$ | The initial coefficient of variation of $x_a$ | 0.125 |
| $CV_{x_h}$ | The initial coefficient of variation of $x_h$ | 0.125 |
| $h^2_{x_a}$ | The narrow-sense heritability of $x_a$ | 0.2 |
| $h^2_{x_h}$ | The narrow sense heritability of $x_h$ | 0.2 |

**S3 Supplemental Figures**

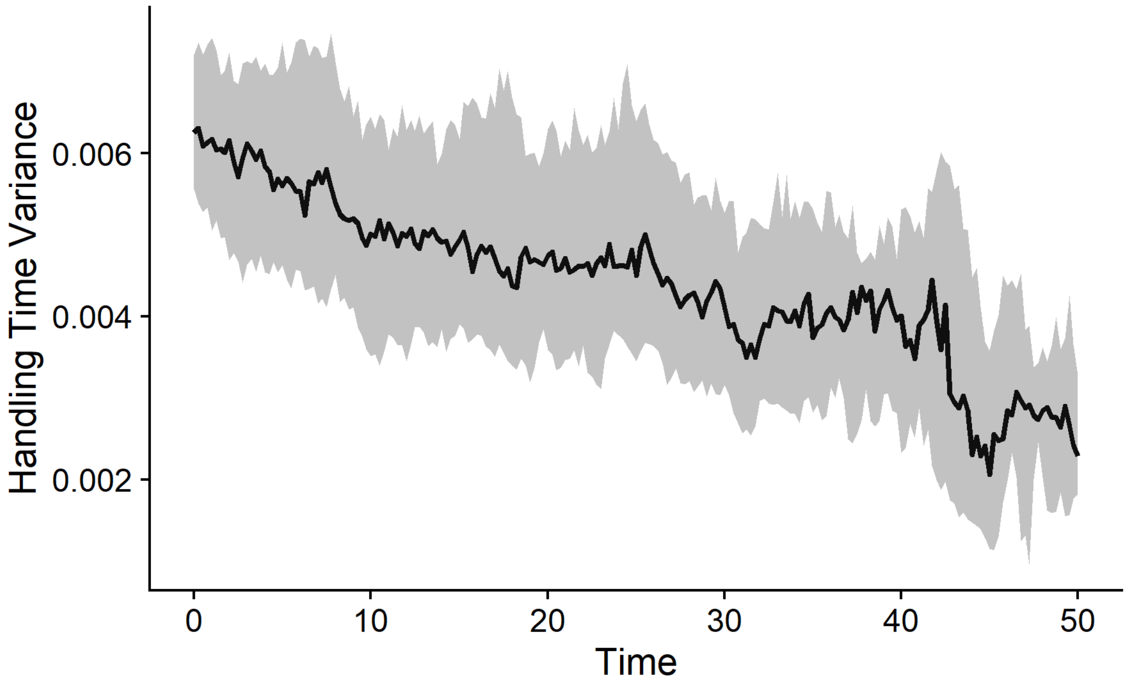


**Figure S3.1.** In the simulations with evolution in the main text with a low carrying capacity, the variance within populations in their handling times decrease over time explaining why evolution of the handling times occurs more slowly than for the quantitative genetics expectation which assumes a constant phenotypic variance in the population. The black lines represent the median trait variance across populations. The gray ribbons represent the 25th and 75th percentiles of variance in the trait within populations.
